# A highly adaptable protocol for mapping spatial features of cellular aggregates in tissues

**DOI:** 10.1101/2024.10.08.617311

**Authors:** Andrew Sawyer, Nick Weingaertner, Ellis Patrick, Carl G Feng

## Abstract

Multiplex imaging technologies have developed rapidly over the past decades. The advancement of multiplex imaging has been driven in part by the recognition the spatial organisation of cells can represent important prognostic biomarkers and that simply studying the composition of cells in diseased tissue is often insufficient. There remains a lack of tools that can perform spatial analysis at the level of cellular aggregates (a common histopathological presentation) such as tumors and granulomas, with most analysis packages focussing on smaller regions of interest and potentially missing patterns in the overall lesion structure and cellular distribution. Here we present a protocol to quantify the cellular structure of entire tissue lesions built around two novel metrics. The Total Cell Preference Index reports whether a lesion tends to change in density in its central vs peripheral areas and can indicate the extent of necrosis across the entire lesion. The Immune Cell Preference Index then reports whether each immune cell-type is located more centrally or peripherally across the entire lesion. The output of both indexes is a single number readout for simple interpretation and visualization, and they can be applied to lesions of any size or shape. Additionally, this protocol can be applied to any slide-scanning multiplexed imaging system, either based on protein and nucleic acid staining. Finally, the protocol uses the open-source software QuPath and can be utilized by researchers with a basic understanding of QuPath with the full protocol able to be applied on pre-generated images within one hour.

## Background

Multiplexed imaging technologies can now perform spatial profiling of dozens of epitopes across millions of cells, all in a single tissue section. These approaches generate enormous amounts of data and can map large, complicated tissue structures such as tumors and granulomas in their entirety within a sectional plane. It is increasingly being shown that the spatial organisation of cells and necrotic tissue in these lesions can represent important prognostic biomarkers, however most analytic tools focus only on cellular composition and / or small regions of interest within the lesions and miss trends at the level of the entire lesion. For example, the extent of leukocyte infiltration into solid tumours has been associated with better prognosis in many cancers (Pagès et al., 2010, Grabovska et al., 2020). Whereas the development of necrosis and lack of cellularity in solid lesion is frequently associated with a poorer prognosis (Liu and Jiao, 2019). As these platforms are able to resolve increasing numbers of cellular markers across larger sections of tissue, they generate ever more complex datasets, and can map tissue structures at high resolution in their entirety (Harms et al., 2023).

Infiltration analysis is a common analysis method to measure the distance of cells from defined tissue lesions (Page et al., 2023, Li et al., 2019). Infiltration analysis requires the lesion border to first be annotated and is then typically performed using a series of software generated bands at fixed distances outward or inward from the lesion border (Figure 1A). The density of the cells of interest is calculated in each of these bands to produce a density histogram to visualise the distribution of a cell of interest within a lesion (Attrill et al., 2022, Tunstall, 2016) (Figure 1B). While infiltration analysis is typically applied to study the relationship between leukocyte populations to a tumour, it can also be used to measure the distance from any cell population to any annotated cellular aggregates. This approach has proven useful for identifying novel biomarkers, it has two key drawbacks. Firstly, it can be confounded when analysing lesions of different size. For example, measuring cells within a 50 μm bandwidth may have different relevance in a lesion of 100 μm diameter compared to a lesion of 1000 μm diameter, making cross-lesion comparison difficult between lesions of different sizes. Furthermore, when the information on multiple bands, cell-types and lesion types are included, the data produced by such analyses is challenging in visualisation, as plotting the total results of an experiment is impractical, requiring an array of heatmaps to visualise the data (Figure 1C, D).

**Figure 1.**
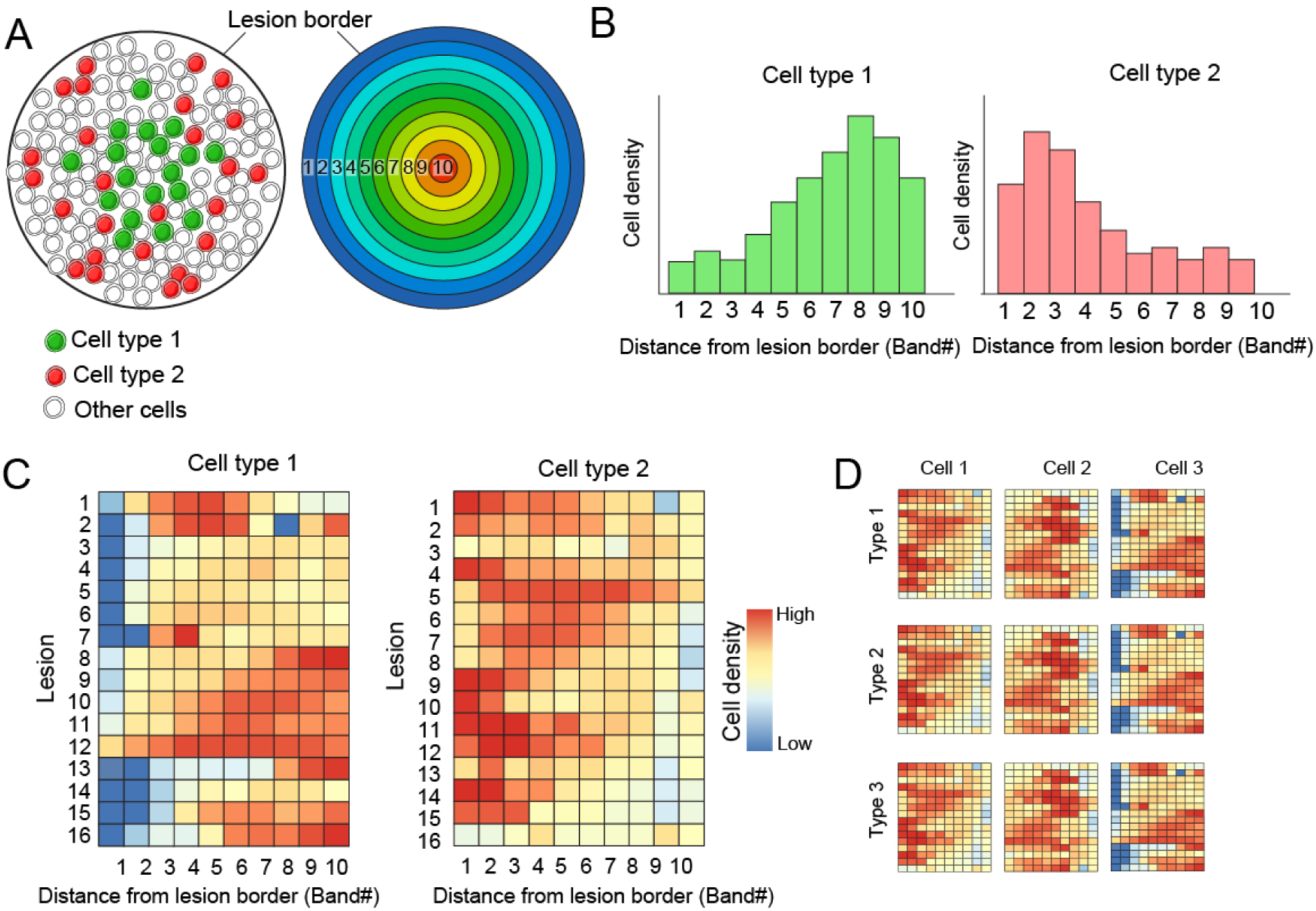
Band-based infiltration analysis is impractical for quantifying intra-lesion cell-type distribution with high lesion throughput. **(A)** Schematic lesion image with cells coloured according to the cell-type (left) Lesion image with bands indicating regions in ten increments from lesion border (right). In both lesion images the lesion border is indicated. **(B)** Example density histograms indicating cell density of two cell types in ten bands from the lesion border in a single image. **(C)** Example heatmaps indicating the cell density of two cell types in ten bands from the lesion border from sixteen lesions. **(D)** Example heatmaps to indicate the cell density of three cell-types and lesions from three lesion types.

Our protocol is designed to analyse multiplex images of entire lesions across the sectional plane. The two tools that this protocol centres on are the total Central Preference Index (tCPI) and the immune Cell Preference Index (immCPI). The tCPI compares measures the density of both the inner and outer 50% of cells in the lesion (relative to the lesion border) and expresses them as a ratio (Figure 2A). This single number output can provide nuanced insight into how necrotic a lesion is and can give an indication of whether a lesion may be losing cells in the centre and tending toward a necrotic phenotype, while not actually being necrotic as of yet (Figure 2B). Additionally, the tCPI tools can be applied to both immunofluorescence (IF) images and haematoxylin and eosin (H&E), as well as any other images that can ascertain the spatial coordinates of each individual cell.

**Figure 2.**
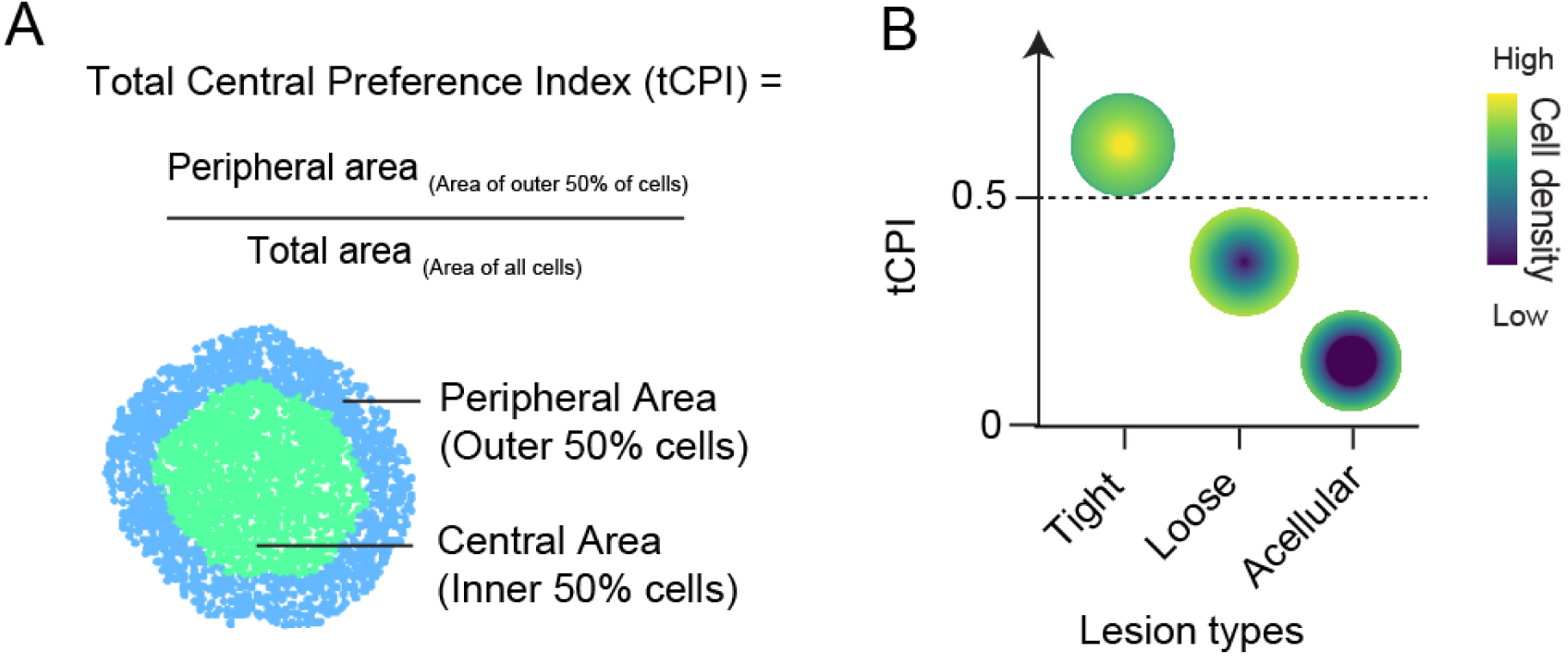
tCPI is a single number measurement for intra - lesion cell localization preference. **(A)** A lesion image with areas coloured according to which cells are closer or further from the lesion border relative to the median distance (top). Equation used to calculate tCPI (bottom). **(B)** Schematic diagram illustrating the intra-lesion cell distribution patterns and their corresponding tCPI.

The immCPI measures the mean distance of all cells of each cell-type (from the lesion border) and compares that with the mean distance of all cells in the lesion (Figure 3A-C). This provides a single number output of whether each cell type tends to localise more centrally or peripherally within the lesion (Figure 3D). A single-number output allows for simple visualisation of the entire datasets with a single bar plot for each cell marker studied (Figure 3E). Importantly both these tools can be applied to lesions of any size and shape. Additionally, this protocol can be applied to any slide-scanning multiplexed imaging system, either based on protein or nucleic acid staining.

**Figure 3.**
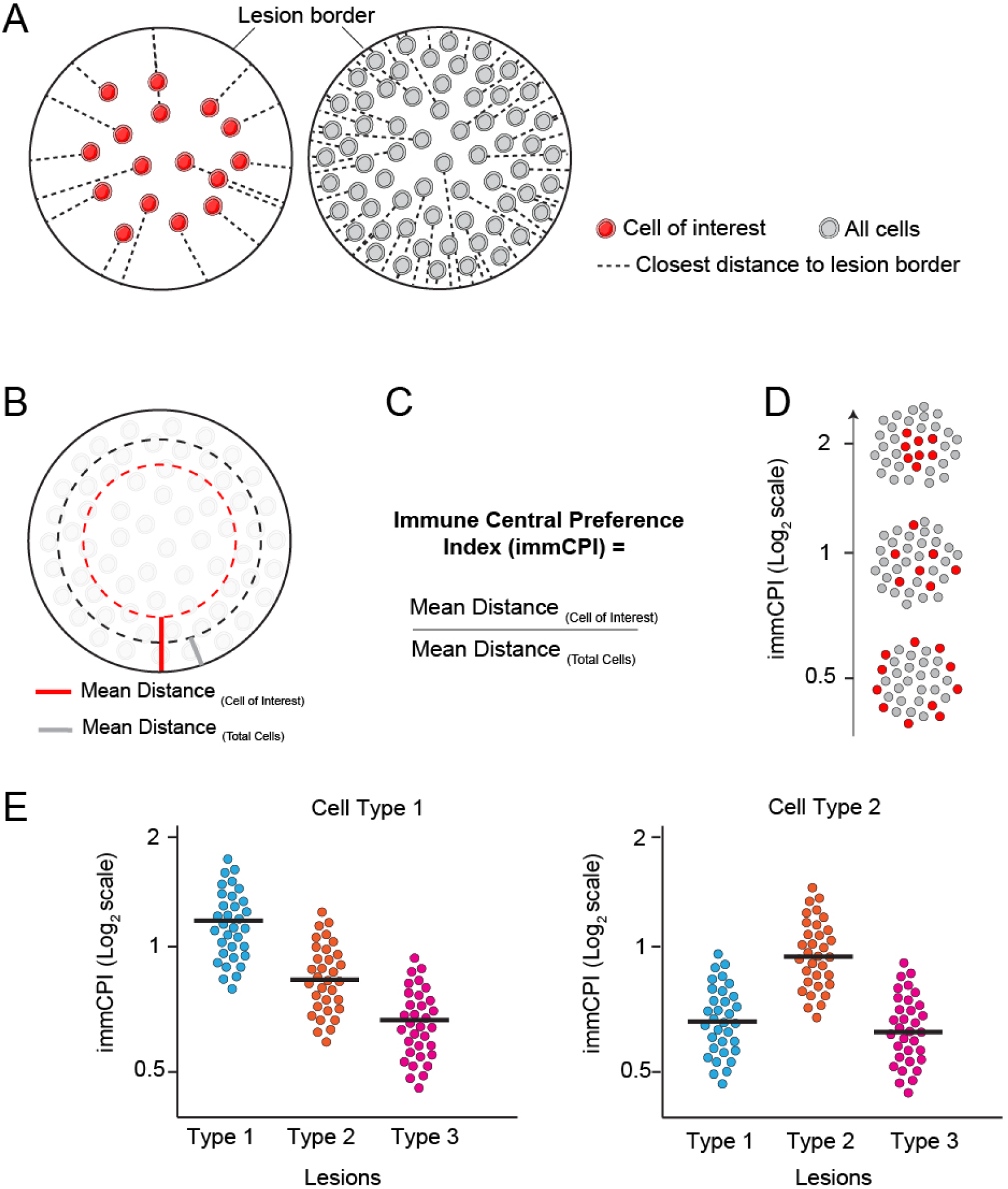
immCPI quantifies distribution of immune-cell populations in any number of lesions and across lesion types. **(A)** A lesion image with only the cell of interest visualized and the closest distance of each cell to the lesion border indicated with dotted line (left). A lesion image with all cells (grey) illustrated and the closest distance of each cell to the lesion border indicated with dotted line (right). **(B)** A schematic lesion image with the mean distance of both the cell of interest and all cells indicated by red and grey bands respectively within the lesion. **(C)** Equation used to calculate immCPI. **(D)** Schematic diagram of lesions with representative immCPI measurements. Red, cell of interest; grey, the rest of the cells **(E)** Example plots comparing the immCPI of two cell types between different lesion types.

This protocol has previously been applied to the study of granulomas in the lungs of tuberculosis patients and revealed a previously unknown level of lesion spatial heterogeneity (Sawyer et al., 2023). The protocol is based on the open-source software QuPath (Bankhead et al., 2017). It can be utilized by researchers with a basic understanding of QuPath with the full protocol able to be applied on pre-generated images within one hour.

## Materials

### Imaging platform

This protocol is applicable to images generated from any multiplexed imaging platform, provided that the area imaged is sufficient to cover the entirety of each tissue lesion examined. Images should be in TIFF format following unmixing and stitching.

### Software

QuPath (version 0.5.1) was used to import TIFF images and

## Procedure

1. Open QuPath and "Create Project"
2. Utilising "Add images", import either whole slide scans or regions containing entire lesions as the appropriate scan type (Brightfield or Immunofluorescence).
3. Utilising the contrast panel change the channel min and max values to improve visibility of stained tissue

### Note

Changing values on the contrast panel will make things easier to see but it will NOT change the underlying image used in analysis. For this reason, ascertaining the correct threshold values is critical at later stages.

4. In the "Annotations" tab, click the three vertical dots or right click the classifications region to expose a menu containing

`Add/Remove->Add Class .` Create a class called "Tissue" and a class called "Lesion".

5. Using the (P)olygon tool within QuPath and a computer mouse or graphics tablet outline the border of the lesion.
6. Select then right click the newly annotated lesion within the viewer and "Set Classification" to "Lesion"
7. Similarly, outline the tissue and assign it the "Tissue" class.
8. Additionally, create a classification for each of your channel names or use the populate from channels function in the right click menu to do this in batch.

### Note

It is possible to use thresholders as an alternative to automatically select the tissue based on the presence of a stain. This works particular well with confluent stains such as eosin. For more information visit the QuPath wiki

9. Open the script editor, denoted by the </> icon in the toolbar
10. Within the file menu, open the tCPI groovy script.
11. Alter the `scanType` variable such that it reflects the nature of the stain (IF or Brightfield)

~~~
// Is it Brightfield or IF
scanType = "IF"
~~~

12. Within the `If` statements for the respective `ScanType`, change the values of the watershed detection to reasonable values for the image being examined.

~~~
if(scanType=="Brightfield"){
runPlugin(’qupath.imagej.detect.cells.WatershedCellDetection’,
’{"detectOD","requestedPixelSizeMicrons":0.4994,"backgroundRadiusMicrons":8.0,"ba
# Change these values where appropriate
…
}
~~~

#### Critical

Correct values for image depend entirely on the image being processed. To trial values, use the GUI controls available under the `Analyse -> Cell Detection -> Cell Detection` menu, noting the values for the script.

Alternative methods to detect cells including Stardist can be utilised instead by replacing the runPlugin command with the Stardist code block.

13. If desired, set `IncludeImmCPI` variable to "Y" in order to run ImmCPI calculation for each marker defined in the header

(Optional).

~~~
// Do you want ImmCPI (Y/N)
def IncludeImmCPI= "Y"
~~~

#### Critical

ImmCPI cannot be calculated based on brightfield imagery as it requires cell type classifications to be made.

14. Set `MarkerNames` variable to include the exact names of the markers used for ImmCPI (Optional).

~~~
// What are the exact names of the marker’s used?
def MarkerNames = ["Opal 690","Opal 650","Opal 570", "Opal 540"];
~~~

15. Define classifiers in `Classify -> Object Classification -> Create Single Measurement Classifier` for each marker utilised. Make sure to name the classification the same as the marker (Optional)

### Note

When creating classifiers, set the channel filter to the appropriate marker, run for all detections in object filter and choose an appropriate measurement type for the cell (nucleus vs cell; mean vs min/max).

16. Create a composite classifier named "All Channels" in `Classify -> Object Classification -> Create Composite Classifier` containing all the markers (Optional).
17. Run the script
18. The tCPI (and optional ImmCPI) will be printed to the console that runs the command, as well as added to the measurement list for the annotation.

~~~
/**
Script to determine the tCPI of an enclosed lesion. Script will detect c Index (tCPI) by Sawyer et al. (2023).
Code to create generate the infiltration annotation is inspired from cod The script requires a lesion annotation with the "Lesion" class and an o Script will by default clear all detections when run. Ensure no detectio To preserve annotations change their class temporarily from "Lesion" so
**/
import qupath.ext.stardist.StarDist2D
import qupath.lib.scripting.QP
import org.locationtech.jts.geom.Geometry
import qupath.lib.common.GeneralTools
import qupath.lib.objects.PathObject
import qupath.lib.objects.PathObjects
import qupath.lib.roi.GeometryTools
import qupath.lib.roi.ROIs
import java.awt.Rectangle
import java.awt.geom.Area
import org.locationtech.jts.precision.GeometryPrecisionReducer
import org.locationtech.jts.geom.PrecisionModel
import qupath.lib.gui.measure.ObservableMeasurementTableData
// ----------------
// Things to modify
// ----------------
// Do you want to keep inner annotation at end of script? Y/N
keepInnerAnnotation = "N"
// Is it Brightfield or IF
scanType = "IF"
// What are the exact names of the marker’s used?
def MarkerNames = ["Opal 690","Opal 650","Opal 570", "Opal 540"];
// Do you want ImmCPI (Y/N)
def IncludeImmCPI= "Y"
// ----------------
// End of Modification Block
// ----------------
~~~

~~~
// Define ImmCPI Calculation
def ImmCpiCalc(ImmuneCellName, anno) {
def ImmuneCellDistance = [];
def RegularCellDistance = [];
getDetectionObjects().each{
if (it.getPathClass()!=null && it.getPathClass().toString().contains(Imm
ImmuneCellDistance.push(it.measurements.get("Signed distance to
}}
getDetectionObjects().each{
RegularCellDistance.push(it.measurements.get("Signed distance to annotat
}
double ImmuneMeanDistance = ImmuneCellDistance.average().toDouble()
double RegularMeanDistance = RegularCellDistance.average().toDouble()
double ImmCpi = ImmuneMeanDistance/RegularMeanDistance
def GranulomaObject = anno
GranulomaObject.measurements.put("ImmCPI("+ImmuneCellName+")",ImmCpi)
print("ImmuneCPI for "+ImmuneCellName+" is: "+ImmCpi)
}
// Define median function not in groovy by standard
def median(data) {
def copy = data.toSorted()
def middle = data.size().intdiv(2)
data.size() %2copy[middle] : (copy[middle-1] + copy[middle])/2
}
// Clear Detections
clearDetections()
selectObjectsByClassification("Lesion")
// Replace the following code blocks with settings for cell detection de
if(scanType=="Brightfield"){
runPlugin(’qupath.imagej.detect.cells.WatershedCellDetection’, ‘{"detect
OD","requestedPixelSizeMicrons":0.4994,"backgroundRadiusMicrons":8.0,"ba
}
if(scanType=="IF"){
runPlugin(’qupath.imagej.detect.cells.WatershedCellDetection’,
‘{"detectionImage":"DAPI","requestedPixelSizeMicrons":0.25,"backgroundRa
if(IncludeImmCPI=="Y"){
runObjectClassifier("All Channels");
}}
//Measure distance from individual cell to lesion border
detectionToAnnotationDistancesSigned(true)
def signedDistances = [];
getDetectionObjects().each {
signedDistances.push(it.measurements.get("Signed distance to annotation
}
medCellDistance = Math.abs(median(signedDistances)).toDouble()
print("Median distance of cells to lesion border is "+medCellDistance.to
PrecisionModel PM = new PrecisionModel(PrecisionModel.FIXED)
//Establish base env Variables
def imageData = getCurrentImageData()
def hierarchy = imageData.getHierarchy()
def server = imageData.getServer()
def cal = server.getPixelCalibration()
if (!cal.hasPixelSizeMicrons()){
print ‘No pixel size calibration detected’
return
}
double expandPixels = medCellDistance / cal.getAveragedPixelSizeMicrons(
def initLesion = getAnnotationObjects().find{it.getPathClass() == getPat
def lesionGeom = getAnnotationObjects().find{it.getPathClass()==getPathC
def plane = ImagePlane.getDefaultPlane()
def tissueGeom = getAnnotationObjects().find{it.getPathClass() == getPat
// Ensure lesion geometry is bound by tissue annotation. This covers cas
lesionIntersectGeom = tissueGeom.intersection(lesionGeom)
lesionROIClean = GeometryTools.geometryToROI(lesionIntersectGeom, plane)
lesionIntersect = PathObjects.createAnnotationObject(lesionROIClean, get
lesionIntersect.setName("Intersected Lesion")
generatedAnnotations = []
generatedAnnotations << lesionIntersect
/**
Following code block adapted from Pete Bankhead’s script at https://pete
**/
// Get the central area
def geomCentral = lesionIntersectGeom.buffer(-expandPixels)
geomCentral = geomCentral.intersection(tissueGeom)
def roiCentral = GeometryTools.geometryToROI(geomCentral, plane)
def annotationCentral = PathObjects.createAnnotationObject(roiCentral)
annotationCentral.setName("Center")
// Get the inner margin area
def geomInner = lesionIntersectGeom
geomInner = geomInner.difference(geomCentral)
geomInner = geomInner.intersection(tissueGeom)
def roiInner = GeometryTools.geometryToROI(geomInner, plane)
def annotationInner = PathObjects.createAnnotationObject(roiInner)
annotationInner.setName("Inner margin")
periph = getPathClass("Periphery")
annotationInner.setPathClass(periph)
addObjects(annotationInner)
/**
--------------
End code block
--------------
~~~

~~~
**/
def regions = []
regions << initLesion
regions << annotationInner
def area = "Area μm^2"
def ob = new ObservableMeasurementTableData()
ob.setImageData(imageData,regions)
double lesionTotalArea = ob.getStringValue(initLesion, area).toDouble()
double peripheralArea = ob.getStringValue(annotationInner, area).toDoubl
double tCPI = peripheralArea/lesionTotalArea
initLesion.measurements.put("tCPI",tCPI)
print("tCPI is "+tCPI)
if(scanType=="IF" && IncludeImmCPI=="Y") {
for (marker in MarkerNames) {
ImmCpiCalc(marker,initLesion)
}
} if(keepInnerAnnotation=="N"){
removeObject(annotationInner, true)
}
~~~

### General notes and troubleshooting

While this protocol can be applied to images generated with any multiplex imaging platform, it is highly advisable to use a tile-based stitched imaging platform, such as the PhenoImager (Akoya) or PhenoCycler Fusion (Akoya), such that the largest possible tissue area can be imaged. This will provide the best chance of capturing lesions in their entirety and has the potential of capturing multiple lesions (potentially dozens from a single slide when lesions are small).

It is important that a reliable method of segmenting individual cells is available as both tCPI and immCPI rely on the spatial locations of each cell centroid. Typically, this will require the addition of a nuclear dye, such as DAPI, to label each cell.

Both the tCPI and immCPI tools also require an annotation of the lesion’s outer border. This can be achieved via multiple means and will differ depending on the disease being studied, which cellular markers are available and whether the image is immunohistochemistry (IHC) or immunofluorescence (IF) based. In tumours, this is often achieved by the identification of a tumour specific marker, such as SOX10 in the case of melanomas (Bahmad et al., 2023). In the case of tuberculosis lesions have been identified through a combination of H&E staining and as high-density aggregates of leukocytes in IF images.

Importantly, while the tCPI tool can be applied to both H&E and IF images, the immCPI tool is only applicable to IF images. tCPI only required cell segmentation to have been performed, which is possible in H&E images by using haematoxylin staining, however immCPI required the cellular identity of each leukocyte population to have been elucidated, which is not possible in H&E images by scripting in QuPath.

### Expected results

The tCPI metric outputs a single number result between 0 and1 for the analysis of each lesion. A result of 0.5 indicates that the inner 50% of cells in a lesion have the same density as the outer 50% of cells from the lesion border and implies that the lesion has a consistent cell density and is unlikely to be necrotic or tending toward a necrotic phenotype. A result above 0.5 indicates that the inner 50% of cells have a higher density of cells compared to the average density of the lesion and implies that the lesion has a tight aggregation of cells in its core. Finally, a score below 0.5 indicates a lesion with a density of the inner 50% of cells less than the average density of the lesion. A score between 0.4 – 0.5 implies that the lesion has a slightly reduced density in its central region and may tend toward a necrotic phenotype, however the lesion may not necessarily having the pathological appearance of a necrotic lesion. A score below 0.4 typically indicates a lesion with pathological necrosis and the closer to 0 the tCPI value, the greater the acellular region within the lesion.

We have previously applied the tCPI tool to the study of 672 tuberculosis lesions from the lungs of 13 patients (Sawyer et al., 2023). We observed at the level of segmented DAPI-positive cells that there appeared to be a high diversity in the tightness and looseness of lesion structures and we identified a spectrum of lesions ranging from tight cellular aggregates to loose potentially pre-necrotic lesions to large necrotizing granulomas (Figure 4A). Using the tCPI tool we were able to confirm that pathologically confirmed necrotizing granulomas had the lowest tCPI score while the pathologically non-necrotizing lesions scores ranged from 0.6 to 0.3 (Figure 4B).

**Figure 4.**
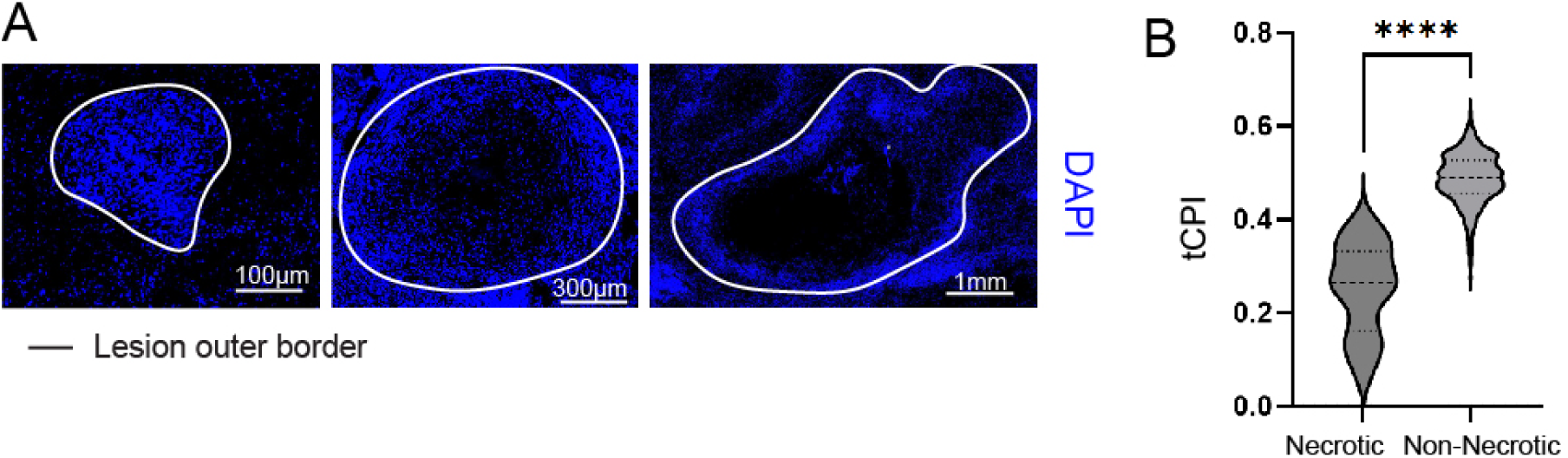
Cells in non-necrotizing lesions show diverse spatial distribution preference. **(A)** Images of DAPI-stained tissue showing variations in intra-lesion cell distribution **(B)** tCPI compared between necrotizing and non-necrotising lesions. The level of statistical significance was determined using a Mann-Whitney test and P <0.05 was considered to indicate statistical significance. **** P < 0.0001.

The immCPI metric outputs a single number result, usually between 0.5 to 2. An immCPI score of 1 for a cell-type of interest within a lesion implies that the mean distance of that cell from the lesion border is equal to the mean distance of all cells in the lesion from the lesion border and indicates that the cell-type is evenly distributed between the lesion periphery and the lesion core. An immCPI value less than 1 implies that the cell of interest is distributed more peripherally in the lesion compared to the mean distance of all cells from the lesion border while an immCPI value greater than 1 implies that a cell of interest is distributed more centrally in the lesion. Importantly, the immCPI tool can provide a comparable output with other lesions regardless of whether any lesions have necrotic cores as immCPI is normalised to all cells in a lesion and automatically excludes necrotic areas from the calculation.

We have previously applied the immCPI tool to the prior cohort of tuberculosis lesions following the identification of 4 major cell populations based on IF staining and 5 classes of lesions based on cell composition and lesion tCPI (Sawyer et al., 2023). We compared the immCPI of each cell type in each lesion across the 5 lesion categories. The immCPI tool revealed that prominent differences between the preference of each cell-type to localise centrally in each lesion class and the immCPI tool allows simple visualisation of all parameters of the analysis (Figure 5).

**Figure 5.**
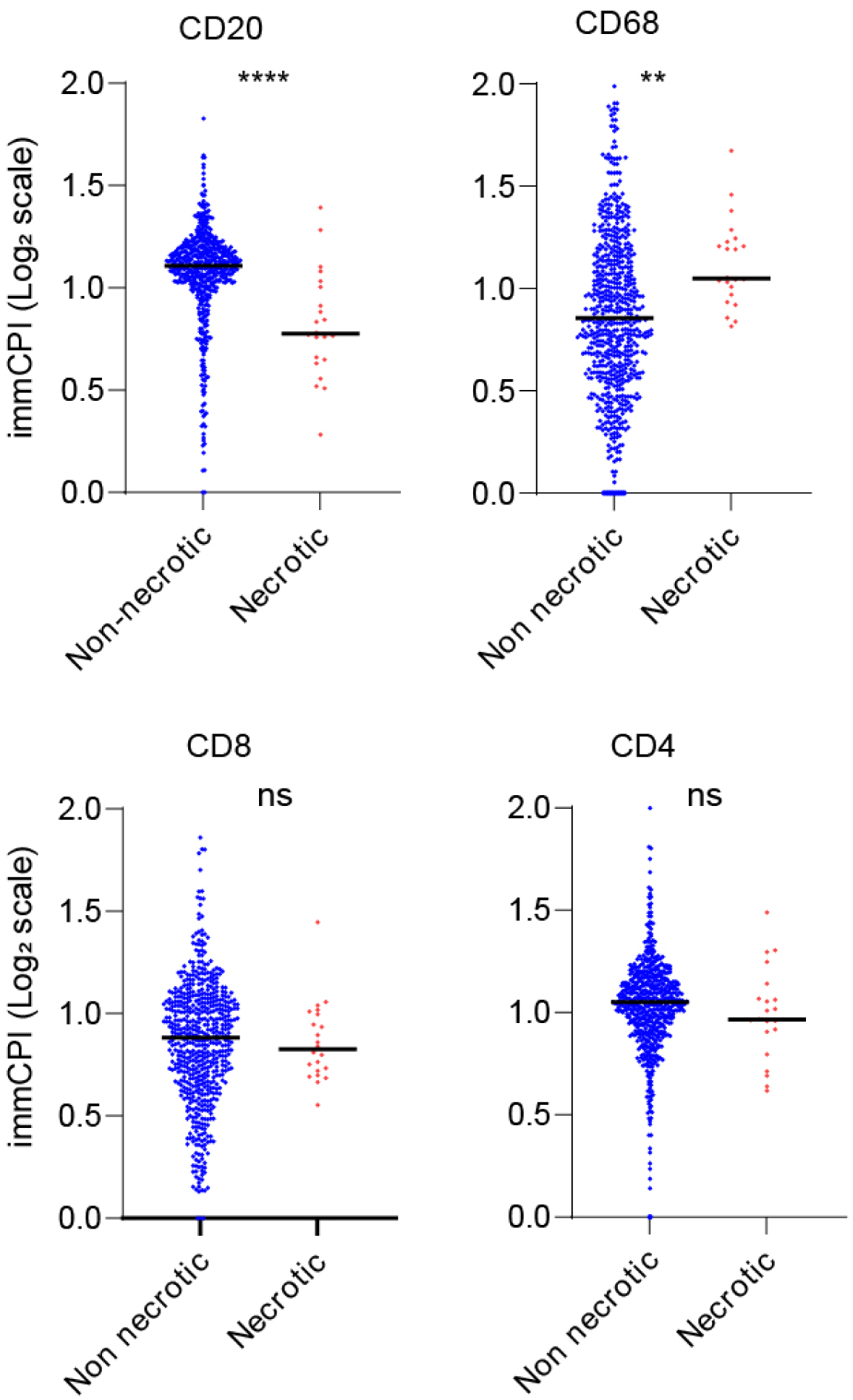
Example showing divergent intra-lesion spatial distribution of CD20+ cells and CD68+ cells between necrotic and non-necrotic tuberculosis granulomas. immCPI of each immune cell type across tuberculosis lesion types. Each symbol represents an individual lesion. Solid black lines indicates group mean. The level of statistical significance is determined with a Mann-Witney test and P < 0.05 was considered to indicate statistical significance. ** P < 0.01. **** P < 0.0001. ns Not significant.

## Conclusion

Here, we provide a workflow for analysis of images from solid tissue lesions generated from any imaging platform. We demonstrate that this workflow can identify the structural characteristics of tuberculosis lesions as an example. Together this workflow will provide a reliable methodology for the analysis of entire cell aggregates and is readily adaptable to future development of imaging platforms to capture tissue lesions in increasing detail.

